# Development of Emerin mRNA Lipid Nanoparticles to Rescue Myogenic Differentiation

**DOI:** 10.1101/2025.06.11.659076

**Authors:** Nicholas Marano, Liza Elif Guner, Rachel S. Riley, James M. Holaska

## Abstract

Emery-Dreifuss muscular dystrophy 1 (EDMD1) arises from mutations in *EMD*. Most EDMD1 patients lack detectable emerin expression. They experience symptoms such as skeletal muscle wasting, joint contractures, and cardiac conduction defects. Currently, physicians rely on treating patient symptoms, without addressing the underlying cause – lack of functional emerin protein. Thus, there is a need for therapeutic approaches that restore emerin protein expression to improve patient outcomes. One way would be to deliver emerin mRNA or protein directly to affected tissues to restore tissue homeostasis. Here, we evaluated the utility of lipid nanoparticles (LNPs) to deliver emerin mRNA to diseased cells. LNPs have been studied for decades and have recently been used clinically for vaccination and treatment of myriad of diseases. Here, we show treatment of emerin-null myogenic progenitors with LNPs encapsulating emerin mRNA causes robust emerin protein expression that persists for at least 4 days. Treatment of differentiating emerin-null myogenic progenitors with 2.5 pg/cell emerin LNPs significantly improved their differentiation. Toxicity profiling of emerin mRNA LNP (EMD-LNP) dosing shows little toxicity at the effective dose. These data support the potential use of EMD-LNPs as a viable treatment option and establishes its utility for studying EDMD pathology.

## Introduction

Emery-Dreifuss Muscular Dystrophy 1 (EDMD) is an X-linked laminopathy caused by mutations in the gene coding for the inner nuclear membrane (INM) protein emerin (Bione et al., 1994). Although emerin is ubiquitously expressed, EDMD1 affects specific tissues, including skeletal muscle and cardiac tissue. Patients experience skeletal muscle wasting, joint contractures, and cardiac defects (Emery and Dreifuss, 1966; Madej-Pilarczyk and Kochanski, 2016). It is common for EDMD1 patients to harbor nonsense mutations in *EMD,* resulting in no detectable emerin expression.

The skeletal muscle pathology of EDMD1 arises, at least in part, from an impaired myogenic differentiation program. Emerin function is critical for myogenic differentiation (Marano and Holaska, 2025). Myogenic progenitors from an emerin-knockout mouse exhibited impaired differentiation with delayed cell cycle withdrawal, belated expression of myogenic markers, and impaired myotube fusion (Iyer et al., 2017; Iyer and Holaska, 2020). Myoblasts from emerin-knockout mice (Melcon et al., 2006) and mouse myoblasts expressing emerin-RNAi (Frock et al., 2006) also showed defective myogenic differentiation. RNA sequencing (RNA-Seq) of differentiating emerin-null H2K myogenic progenitors showed the temporal expression of cell cycle and myogenic lineage genes was perturbed (Iyer and Holaska, 2020). This is likely due to loss of key emerin interactions with various binding partners, including histone modifying machinery such as HDAC3 (Demmerle et al., 2012), G9a and EZH2 (Marano and Holaska, 2022), transcription regulatory proteins (Holaska et al., 2003; Haraguchi et al., 2004; Holaska and Wilson, 2007; Lee et al., 2017) and proteins that maintain nuclear architecture (Holaska et al., 2004; Haque et al., 2010; Fernandez et al., 2022).

Clinical interventions for EDMD1 patients rely on treating symptoms. For example, joint contractures can be treated with physical therapy or surgical intervention to improve joint mobility and reduce pain (Bonne et al., 1993). Cardiac tissue is also affected in EDMD1, and treatments include angiotensin converting enzyme (ACE) inhibitors or angiotensin receptor blockers (ARBs) to assist with arrythmias (Heller et al., 2020). As the disease progresses, patients often require a pacemaker to maintain proper cardiac function (Bialer et al., 1991; Steckiewicz et al., 2016)

Currently, gene therapy treatment options for EDMD1 are lacking. To address this problem, we utilized lipid nanoparticle (LNP) technology to deliver emerin mRNA to cells (Hou et al., 2021). LNPs have been studied extensively for nucleic acid delivery both *in vitro* and *in vivo* in cancer (Chen et al., 2022; Lei et al., 2024), viral indications (Polack et al., 2020; Baden et al., 2021), genetic diseases (Adams et al., 2018), pregnancy (Young et al., 2024; Hofbauer et al., 2025), and more, with several platforms having gained FDA approval (Adams et al., 2018; Polack et al., 2020; Baden et al., 2021). However, LNPs have not been developed or tested as a therapeutic strategy for EDMD1. These delivery vehicles offer protection over the nucleic acid cargo, intracellular uptake, and cytosolic nucleic acid delivery, overcoming many nucleic acid delivery challenges (Riley et al., 2019). Moreover, these platforms are highly safe and are well tolerated in clinical studies. Therefore, LNP-mediated mRNA delivery has emerged as a promising avenue for expression of transient proteins. Here, we encapsulated human emerin mRNA within LNPs (EMD-LNPs) and show that this platform delivers robust emerin expression in emerin-null myogenic progenitors while showing minimal toxicity. Importantly, EMD-LNPs rescue emerin-null myogenic differentiation to wildtype levels.

## Materials and Methods

### Cell Culture

Myogenic progenitors from H2K wildtype and EMD^-/y^ mice were obtained from Tatiana Cohen and Terence Partridge (Children’s National Medical Center, Washington DC, United States). Proliferating myogenic progenitors were collected and counted by a Countess® II FL Automated Cell Counter (ThermoFisher Scientific). Cells were plated at 650 cells/cm^2^ and incubated in proliferative media consisting of high-glucose DMEM (ThermoFisher Scientific) supplemented with 2% chick embryo extract (Accurate Chemical; #MDL-004-E), 20% heat-inactivated FBS (ThermoFisher Scientific; #16000048), 1% penicillin-streptomycin (ThermoFisher Scientific), 2% L-glutamine (ThermoFisher Scientific) and 20 units/mL γ-interferon (MilliporeSigma) at 33°C and 10% CO_2_. For differentiation experiments, cells were plated at 25,000 cells/cm^2^ in proliferative media at 33°C and 10% CO_2_ overnight. Cells were washed once with phosphate-buffered saline (PBS) and incubated with differentiation media consisting of high-glucose DMEM, 2% horse serum, and 1% penicillin/streptomycin at 37°C and 5% CO_2_.

### Formulation of EMD-LNPs

C12-200 was purchased from MedChem Express (Monmouth Junction, NJ) and all other lipid components, including DOPE, cholesterol, and DMPE-PEG2000, were purchased from Avanti Polar Lipids Inc. (Birmingham, AL). Codon optimized luciferase and emerin mRNA (NM_000117.3) were purchased from the Engineered mRNA and Targeted Nanomedicine core facility at the University of Pennsylvania. The mRNA was designed with 1-methylpseudouridine modifications and is co-transcriptionally capped and purified using cellulose-based chromatography. LNPs were formulated by micropipette mixing of an ethanol phase, containing the lipid components, and an aqueous phase, containing the mRNA and citrate buffer (pH 3), at a 1:3 volumetric ratio. The formulation contained a molar ratio of 35% ionizable lipid, 10% phospholipid, 1.5% DMPE-PEG, and 53.5% cholesterol. The mRNA:ionizable lipid weight ratio was 1:10. After synthesis, the LNPs were dialyzed against 1X PBS for 2 hours and sterile filtered using 0.2 µm syringe filters.

### Characterization of EMD-LNPs

Dynamic Light Scattering (DLS) measurements, including hydrodynamic diameter and polydispersity index, were taken with a Malvern Panalytical Zetasizer Nano ZS (Malvern, UK). LNP samples were diluted 1:100 with PBS in cuvettes before taking DLS measurements. LNP encapsulation efficiencies were calculated with the Promega QuantiFluor® RNA System kit (Madison, WI). Briefly, LNPs were diluted in 1X TE buffer to a 1:100 ratio in two tubes, with one tube containing 1% Triton X-100. Samples were then heated to 37°C under agitation (300 rpm) for 10 minutes. A standard curve of mRNA was prepared to quantify encapsulation within LNPs per manufacturer instructions. Once cooled to room temperature, LNP samples and standards were plated in a black 96-well plate in triplicate. Fluorescent dye was then added to each well and fluorescent signal was read on a plate reader (excitation: 494 nm, emission: 540 nm).

### Differentiation Assay

Wildtype and EMD^-/y^ myogenic progenitors were counted and plated in 12-well dishes (Falcon) at a density of 25,000 cells/cm^2^ and grown in proliferative conditions for 2 hours. Cells were then dosed at a concentration of 2.5 pg/cell and incubated overnight. Cells were then washed and induced to differentiate (t=0 hr.) by replacing with differentiation media and grown in differentiation conditions. Plates were grown for 72 hours and samples were collected every 24 hours. The ClickIt EdU reaction kit (ThermoFisher Scientific #C10640) was used to determine cell cycle withdrawal. After 22 hours in differentiation conditions, EdU was added to cells and allowed to incubate for two hours. Samples were washed once with PBS, fixed with 3.7% formaldehyde for 15 minutes, permeabilized in 0.2% triton X-100 for 20 minutes, and stained for EdU incorporation per manufacturer instructions. To determine myogenic differentiation index and fusion index, cells were grown for 48 or 72 hours, respectively, in differentiation conditions then fixed and permeabilized as previously described. Cells were blocked with 3% BSA in PBS at room temperature for at least 1 hour with shaking. Cells were washed and incubated with myosin heavy chain (MyHC) antibody (1:20; Santa Cruz Biotechnology; #SC-376157) for 2 hours at room temperature with shaking. Cells were then washed and incubated with secondary antibody anti-mouse Alexa-594 (Invitrogen #A-11032). Nuclei were stained with 0.2 µg/mL DAPI in PBS for 5 minutes at room temperature with shaking. Images were taken with a 40X objective (N.A.=0.65) equipped on an Evos FL Auto 2 fluorescent microscope. Cell cycle withdrawal was calculated by determining the ratio of EdU-positive (Alexa-647) nuclei to total number of nuclei (DAPI). Differentiation index was calculated by determining the ratio of MyHC-positive (Alexa-594) cells to total number of nuclei (DAPI). Fusion index was calculated by determining the ratio of nuclei (≥3) contained in a shared MyHC-positive cytoplasm to total number of nuclei (DAPI). Images were analyzed and counted using imageJ.

### Western Blotting

Whole cell lysates were made using NuPage™ LDS sample buffer (Invitrogen #NP007). Samples were heated at 75°C for 15 minutes on a heating block prior to electrophoresis. Samples were then separated on a NuPage bis-tris gel (Invitrogen #NP0321BOX) at constant voltage for one hour. Samples were then horizontally transferred to a nitrocellulose membrane under constant 250mA current for 75 minutes on ice. Membranes were stained with ponceau S to verify transfer efficiency. Membranes were washed with PBS containing 0.1% triton X-100 (PBST) and blocked with 5% milk (Giant Food Stores) diluted into PBST for one hour at room temperature or overnight at 4°C. Membranes were stained with primary antibodies against emerin (1:3,000; ProteinTech #10351-1-AP), *γ*-tubulin (1:10,000; Invitrogen #), H3K9me2 (1:1,500; Active Motif #39239), or H4K5ac (1:1,500; Sigma-Aldrich #07-327) diluted in 0.5% milk in PBST. All primary antibodies were incubated for one hour at room temperature with shaking or overnight at 4°C with shaking. Membranes were washed at least 3 times for 5 minutes in PBST and incubated with HRP-conjugated secondary antibodies against mouse or rabbit for one hour at room temperature. Membranes were then washed at least 3 times in PBST and incubated for one minute in SuperSignal™ West PicoPlus chemiluminescent substrate (Life Technologies #34580) before visualizing on a Li-Cor Odyssey Fc imager. Densitometry analysis was performed in Li-Cor Image Studio software. Bands were normalized to *γ*-tubulin as a loading control. All washing and incubations steps were performed with moderate shaking.

### Immunofluorescence

Myogenic progenitors were grown in 48-well dishes and dosed with EMD-LNPs or PBS. Cells were fixed in 3.7% formaldehyde for 15 minutes. Cells were washed and permeabilized with 0.2% triton X-100 for 20 minutes. Cells were washed and blocked with 3% BSA in PBS for 1 hour. Primary emerin antibody (1:500; ProteinTech #10351-1-AP) was diluted in blocking buffer and incubated for 2 hours at room temperature or overnight at 4°C. Cells were washed 3 times with PBS for 5 minutes and allowed to incubate with Alexa Fluor™-conjugated secondary antibodies against rabbit (Invitrogen) for one hour at room temperature shielded from light. All washing and incubations steps were performed with moderate shaking.

### Cell Viability Assay

EMD^-/y^ myogenic progenitors were plated in a 96-well opaque-walled dish with clear bottoms at a concentration of 1,000 cells/well. The cells were allowed to adhere to the plate for 2 hours in proliferative conditions. Cells were then dosed with PBS, luciferase mRNA LNPs (LUC-LNPs), or EMD-LNPs at varying concentrations and grown overnight in proliferative conditions. Cells were washed twice with PBS and replaced with proliferative media (no LNP) containing PrestoBlue reagent and allowed to incubate for 1 hour at 33°C and 10% CO_2_. The plate was then analyzed for fluorescence on a Flex Station 3 plate reader and background of PrestoBlue media-only was subtracted from each readout. Following fluorescence detection, cells were washed and replaced with proliferative media and grown 24 hours in proliferative conditions. This process was repeated for 2 additional days.

## Results

### LNP synthesis and characterization

In this study, LNPs were comprised of 35% C12-200 ionizable lipid, 10% DOPE, 1.5% DMPE-PEG, and 53.5% cholesterol, based on our prior studies that utilized this formulation for mRNA delivery (Young et al., 2024; Hofbauer et al., 2025). Here, we used the ionizable lipid C12-200 because it has been extensively evaluated for nucleic acid delivery to a variety of tissues and cell types (Love et al., 2010; Kauffman et al., 2015; da Silva et al., 2024; Young et al., 2024). The ionizable lipid component is pH-responsive and becomes protonated in acidic environments. This interaction allows for endosomal escape once taken up by cells and is the main driver of RNA encapsulation within LNPs (Riley et al., 2021). Either luciferase or emerin mRNA was encapsulated within LNPs for experimental analysis. On average, LNPs with luciferase mRNA were 115.15 nm in diameter and had 36.58 (μg/mL) mRNA encapsulation (Fig. S1A,B). Similarly, LNPs with emerin mRNA were 121.36 nm in diameter and had 33.39 (μg/mL) mRNA encapsulation (Fig. S1A,B). All LNPs were highly monodisperse with polydispersity indices below 0.2 (Fig. S1C).

### Lipid nanoparticles efficiently deliver emerin mRNA to H2K myogenic progenitors

After incubating EMD-LNPs (2.5 pg/cell) with emerin-null myogenic progenitors for 24 hours, >98% of nuclei expressed emerin (Figure 1A). >95% of nuclei retained emerin expression after 96 hours (Figure 1B). Emerin protein expression was measured by western blotting, which showed increased human emerin expression following LNP treatment. It is important to note that the human emerin produced by the mRNA ran at a lower molecular weight than wildtype mouse emerin (Figure 1C). This is because human emerin is smaller than the mouse isoform. 24 hours after EMD-LNP treatment of emerin-null progenitors, emerin levels were 7.72-fold (±1.98) greater than endogenous emerin in wildtype myogenic progenitors (Figure 1C,D). Western blotting every day for 4 days (Figure 1C,D), showed emerin protein expression decreased over time. At 48, 72, and 96 hours after EMD-LNP dosing, emerin expression in emerin-null myogenic progenitors was 4.31 (±1.02)-fold, 0.98 (±0.29)-fold, and 0.44 (±0.03)-fold that of wildtype emerin levels, respectively (Figure 1 C,D).

**Figure 1.**
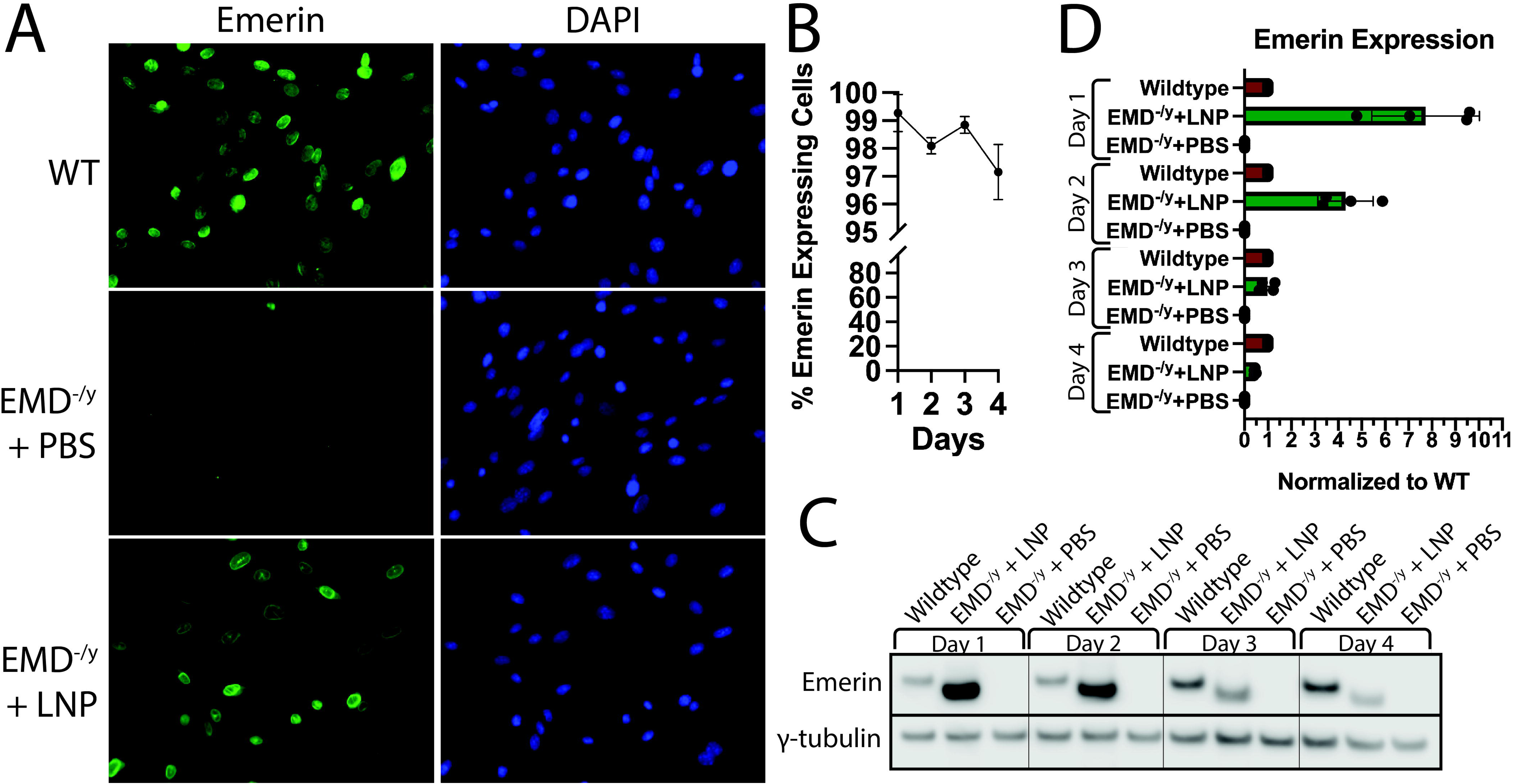
EMD-LNPs efficiently deliver cargo *in vitro.* A, Emerin-null myogenic progenitors dosed with 2.5 pg/cell EMD-LNPs (EMD^-/y^ + LNP) or PBS (EMD^-/y^ + PBS) for 22 hours. Immunofluorescence microscopy was performed using emerin antibodies (green). DAPI (blue), DNA. B, Most EMD-LNP-treated emerin-null progenitors retain emerin protein expression after 4 days. C, Western blotting was performed on wildtype (WT) or emerin-null proliferative myogenic progenitors treated with EMD-LNPs or (EMD^-/y^ + LNP) or PBS (EMD^-/y^ + PBS) over four days. D, Quantitation of emerin protein expression normalized to endogenous emerin in wildtype progenitors. (N=4)

### LNPs yield no toxicity at effective treatment doses

We next determined the cytotoxic profile of LNPs containing emerin mRNA or luciferase mRNA (LUC-LNP). We utilized the PrestoBlue cell viability reagent, which measures cellular metabolism by the conversion of a non-fluorescent substrate (resazurin) into a fluorescent product (resorufin). Cells were dosed with increasing concentrations of emerin or luciferase mRNA LNP (0 – 60 pg/cell) or PBS (equivalent volume). Cells were incubated with LNPs or control for 22 hours and PrestoBlue cell viability assays were performed over 3 days (Figure 2; S2). Dosing of EMD-LNPs and LUC-LNPs <15 pg/cell were non-toxic (Figure 2, S2), which is well above the effective doses used for these studies (2.5 pg/cell). However, EMD-LNP and LUC-LNP dosing at ≥15 pg/cell had significantly reduced signal 24 hours after treatment, compared to PBS control (Figure 2A). EMD-LNP doses at 15, 30, and 60 pg/cell reduced signal by 58.7% ± 4.1%, 81.2% ± 5.0% and 93.9% ± 4.2%, respectively, when compared to PBS controls. While viability was significantly reduced at these doses, only at 60 pg mRNA/cell were cells non-viable, as observed by PrestoBlue (Figure 2A) and by brightfield microscopy (not shown). To monitor LNP cytotoxicity over time, these cells were washed with PBS and replaced with proliferative media overnight. This process was repeated every 24 hours for 3 days (Figure 2, S2). Significant decreases in fluorescence were also observed for both emerin and luciferase mRNA LNPs at concentrations ≥15 pg/cell at 48 hours (Figure 2B) and 72 hours (Figure 2C), indicative of cytotoxicity. 48 hours after LNP dosing, cells dosed with 15, 30, and 60 pg/cell showed 44.1% ± 7.1%, 79.2% ± 1.3%, and 96.8% ± 0.2% reductions in signal when compared to PBS controls, respectively (Figure 2B). Similarly at 72 hours, 15, 30, and 60 pg/cell doses led to 41.4% ± 5.7%, 80.2% ± 0.3%, and 97.8% ± 0.9% reductions in signal compared to PBS controls, respectively (Figure 2C).

**Figure 2.**
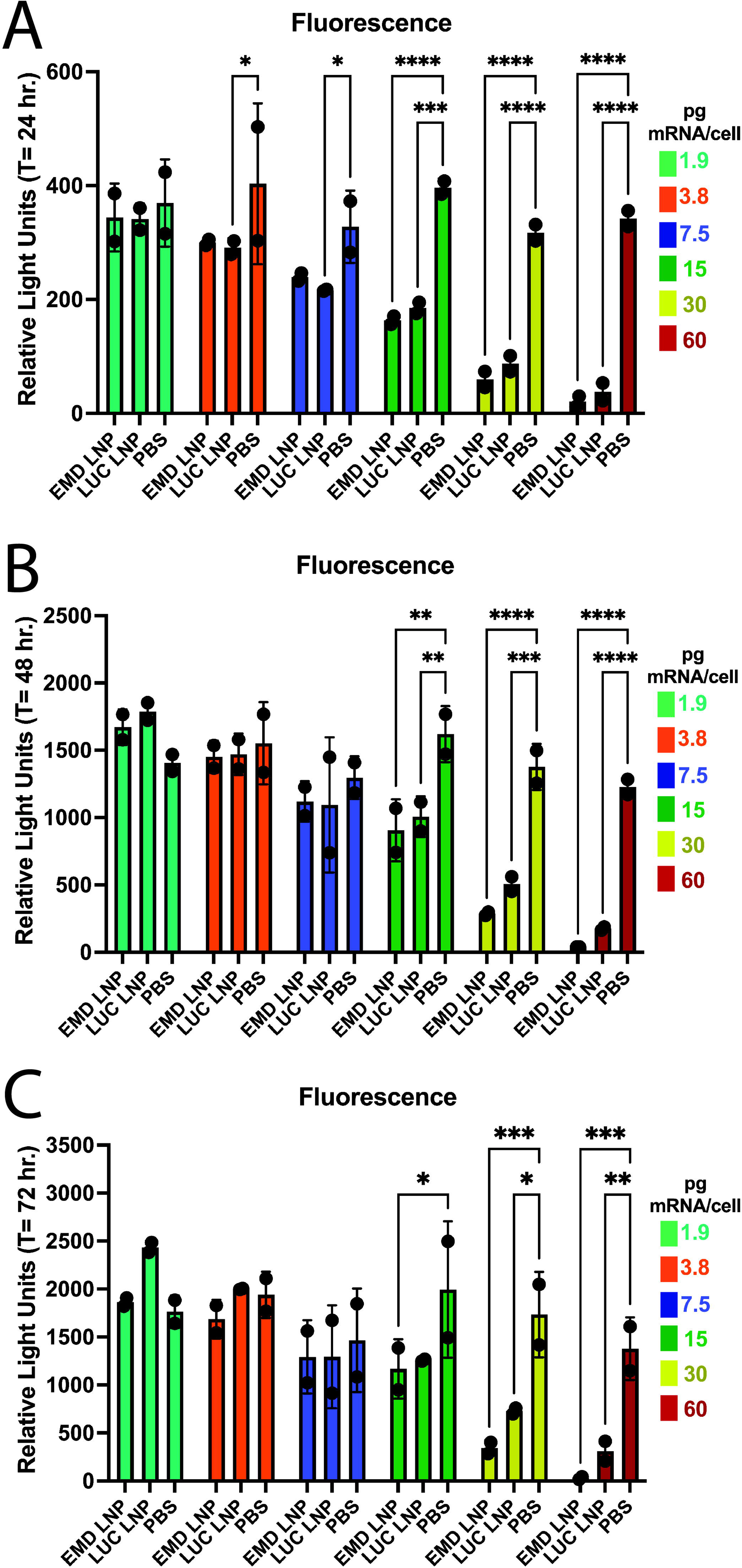
EMD-LNPs become cytotoxic at ≥15 pg/cell. PrestoBlue cell viability assays were used to measure cellular metabolism in EMD^-/y^ + PBS (PBS), EMD^-/y^ + EMD-LNP (EMD LNP), or EMD^-/y^ + luciferase mRNA LNP (LUC LNP) proliferating myogenic progenitors with increasing concentrations of LNPs. Measurements were recorded every 24 hours for 72 hours. A, 24 hours after LNP incubation. B, 48 hours after LNP incubation. C, 72 hours after LNP incubation (N=2) (* p≤0.05; ** p≤0.01; *** p≤0.001)

### EMD-LNPs rescue differentiation of emerin-null myogenic progenitors

We showed that stable transduction of wildtype emerin into emerin-null H2K myogenic progenitors rescued cell cycle withdrawal, differentiation index, and fusion index to wildtype levels (Iyer et al., 2017; Marano and Holaska, 2022). Therefore, we wanted to assess the ability of EMD-LNPs to rescue differentiation, a requirement for therapeutic use. We performed differentiation assays (Figure 3A) on emerin-null myogenic progenitors treated with emerin LNPs or PBS, as previously described (Iyer et al., 2017; Iyer and Holaska, 2020; Marano and Holaska, 2022). We analyzed cell cycle withdrawal using EdU incorporation to determine the proportion of actively dividing or recently divided cells (Figure 3B). Wildtype H2K myogenic progenitors rapidly exit the cell cycle with 14.9% ± 2.0% still cycling at 24 hours post-differentiation induction (Figure 3C). Alternatively, 20.3% ± 1.0% of emerin-null myogenic progenitor nuclei were positive for EdU after PBS treatment, showing impaired cell cycle exit when compared to wildtype (Figure 3B,C). Treatment of emerin-null progenitors with emerin LNPs rescued cell cycle exit, as 16.6% ± 2.1% of these nuclei were EdU-positive (Figure 3 B,C). 48 hours after differentiation induction, wildtype or emerin-null cells dosed with 2.5 pg/cell EMD-LNP or PBS were fixed and stained for myosin heavy chain (MyHC) to measure the differentiation index (Figure 3D,E). The differentiation index is defined as the number of cells with MyHC-positive cytoplasm compared to the total number of cells (VanGenderen et al., 2022). As expected, 54.6% ± 1.7% of emerin-null progenitors dosed with PBS expressed MyHC compared to 66.1% ± 1.8% of wildtype progenitors (Figure 3 D,E), showing impaired differentiation. Significantly more emerin LNP-dosed emerin-null progenitors expressed MyHC (62.7% ± 1.3%; Figure 3 D,E) demonstrating a rescue of differentiation by EMD-LNPs.

**Figure 3.**
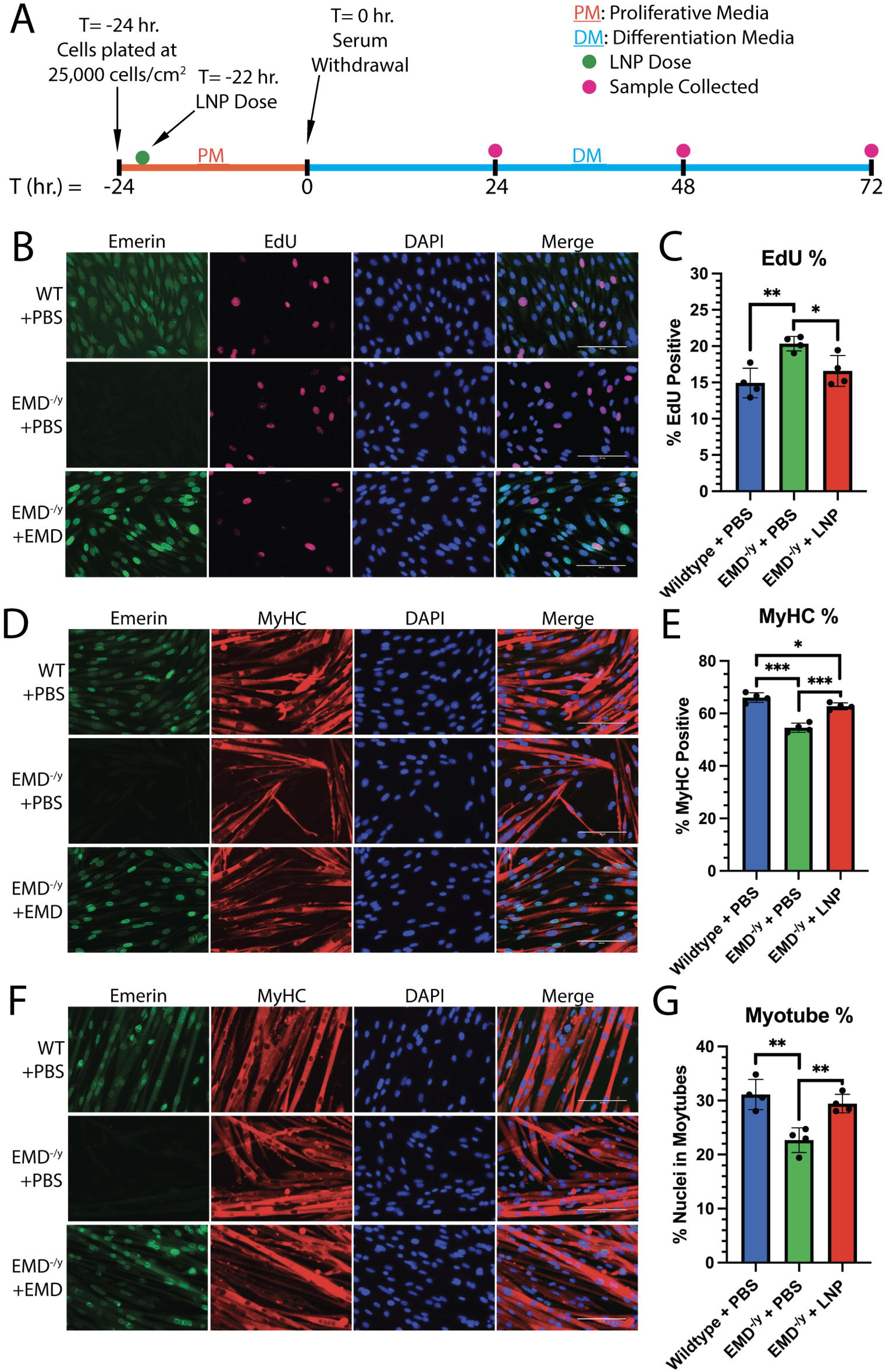
EMD-LNPs rescue differentiation of emerin-null myogenic progenitors. A, Line graph showing the timing of LNP treatment (green dot), differentiation induction, and sample collection (pink dot). Wildtype myogenic progenitors treated with PBS (WT + PBS), emerin-null progenitors treated with PBS (EMD^-/y^ + PBS), or emerin-null progenitors treated with 2.5 pg/cell of EMD-LNPs (EMD^-/y^ + LNP) were incubated for 22 hours and induced to differentiate. B, After 24 hours the cells were incubated with EdU for 2 hours. The cells were fixed and incubated with emerin antibodies. EdU, red; emerin, green; DAPI, DNA (blue). C, Quantification of EdU-positive nuclei. D,F Immunofluorescence microscopy was done on after 48 hours (D) or 72 hours (F), to detect MyHC (red) or emerin (green). DAPI, DNA (blue). E, Differentiation index was quantified by determining the percentage of MyHC-positive cells. F, Fusion index represents the percentage of nuclei present in a shared (≥3 nuclei/cell) MyHC-positive cytoplasm. (N=4; ≥100 nuclei per biological replicate) (* p≤0.05; ** p≤0.01; *** p≤0.001)

The fusion index was measured 72 hours after differentiation induction to assess myoblast fusion. The fusion index is defined as the number of nuclei within myotubes (cells with ≥3 nuclei in a shared MyHC-positive cytoplasm) divided by the total number of nuclei (VanGenderen et al., 2022). 31.1% ± 2.8% of wildtype nuclei were present in myotubes (Figure 3 F,G). Emerin-null progenitors had a significantly reduced fusion index with 22.7% ± 2.3% of nuclei present in myotubes (Figure 3F,G). Treatment with emerin LNPs increased the fusion index to 29.4% ± 1.7% (Figure 3F,G). Therefore, our data suggests that treatment of emerin-null myogenic progenitors with 2.5 ng/cell EMD-LNPs rescues myogenic differentiation.

### EMD-LNPs increase repressive histone modification levels

Our previous data suggests that the coordinated temporal expression of differentiation genes is dysregulated in emerin-null progenitors (Koch and Holaska, 2012; Iyer et al., 2017). Previous studies also suggest this may result from defective genomic reorganization of these genomic loci during differentiation (Demmerle et al., 2013). Thus, we tested if emerin LNPs affect genomic organization dynamics by altering histone modifications, as was seen for stable expression of wildtype emerin in emerin-null myogenic progenitors (Demmerle et al., 2012). We examined H4K5ac and H3K9me2, since they are implicated in repressive genomic organization at the nuclear envelope. We dosed proliferative emerin-null myogenic progenitors with 2.5 pg/cell for 22 hours and measured the levels of H3K9me2 and H4K5ac (Figure 4A). EMD-LNP treatment increased H3K9me2 levels by 26.5% ± 15.5% compared to wildtype progenitors (Figure 4B). Compared to emerin-null PBS controls, LNP-treated emerin-null myogenic progenitors had a 20.5% ± 10.0% reduction in H4K5ac levels (Figure 4C).

**Figure 4.**
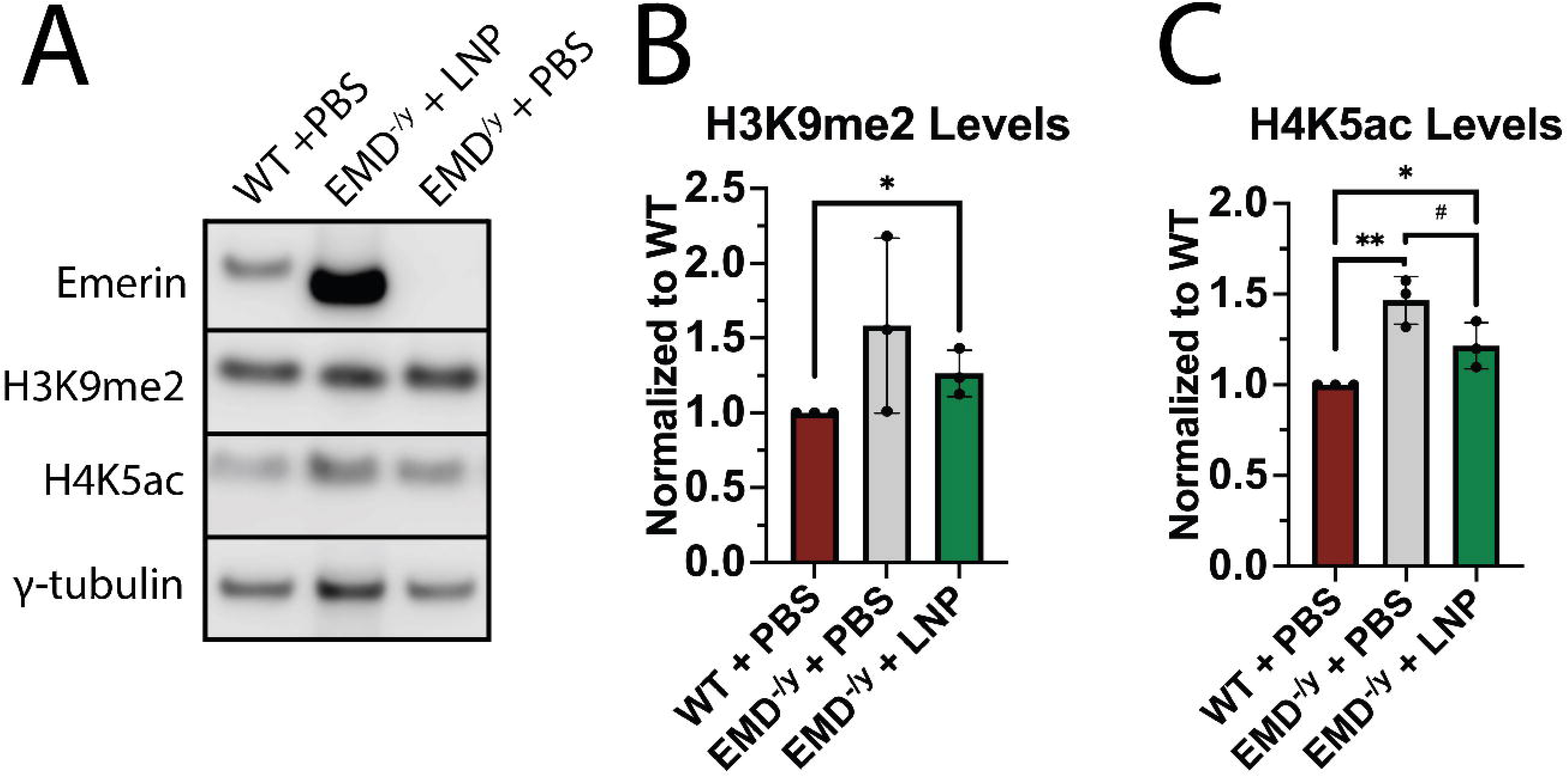
EMD-LNPs partially rescue H4K5ac and H3K9me2 levels in emerin-null myogenic progenitors. Whole cell lysates were collected from wildtype cells treated with PBS (WT + PBS), emerin-null cells treated with 2.5 pg/cell of EMD-LNPs (EMD^-/y^ + LNP), and emerin-null cells treated with PBS (EMD^-/y^ + PBS) 24 hours after treatment. A, Western blotting for emerin, H3K9me2, H4K5ac, and *γ*-tubulin. B,C Quantification of H3K9me2 and H4K5ac western blots. Levels of each protein were normalized to *γ*-tubulin and then normalized to wildtype cells. (N=3) (# p≤0.08; * p≤0.05; ** p≤0.01)

## Discussion

Here we showed EMD-LNPs can successfully deliver mRNA cargo to myogenic progenitors. Upon treatment, emerin is expressed at high levels and remains for at least 4 days. We also established the dose at which EMD-LNPs become cytotoxic to be ≥15 pg/cell – much greater than the safe dose of 2.5 pg/cell used in this analysis. We have previously shown the expression of wildtype emerin in emerin-null myogenic progenitors rescues differentiation to wildtype levels (Iyer and Holaska, 2020; Marano and Holaska, 2022). Here we show that expression of wildtype emerin in emerin-null cells can be accomplished by addition of EMD-LNPs and that this rescues emerin-null myogenic progenitor differentiation. The skeletal muscle pathology of EDMD1 arises, at least in part, from poor myogenic differentiation. Thus, we predict EMD-LNP treatment would ameliorate the skeletal muscle symptoms of EDMD1 patients.

Emerin interaction with HDAC3 has been shown to activate catalytic activity, leading to reduced H4K5ac levels (Demmerle et al., 2012). Additionally, targeting H4K5ac levels by an HDAC3 activator (theophylline) or inhibitors of H4K5-specific histone acetyltransferases rescued differentiation and fusion indices of emerin-null myogenic progenitors (Collins et al., 2017). Here we also observed significant reductions in H4K5ac levels of emerin-null progenitors dosed with EMD-LNPs. It is possible the changes in H4K5ac levels are a direct result of the emerin-HDAC3 interaction and activation of HDAC3 catalytic activity. HDAC3 activation in emerin-null myogenic progenitors leads to the reorganization of specific stem cell and myogenic gene loci to resemble wildtype progenitor genomic organization (Demmerle et al., 2013). Therefore, rescue of differentiation by expression of emerin in emerin-null myogenic progenitors may be due to emerin-HDAC3 interaction and proper genomic organization or epigenetic response that leads to proper myogenesis.

This data supports the use of emerin LNP as a potential therapeutic intervention for EDMD1 that requires more study to ascertain its effectiveness and feasibility. Other gene therapy interventions to treat muscular dystrophies are currently in development, which include those that utilize adeno-associated viruses (AAV) targeted to muscle cells (myo-AAV) to deliver and integrate exogenous DNA into cells (Tabebordbar et al., 2021; Ji et al., 2024). LNP technology may be advantageous over Myo-AAVs for a couple of reasons. First, EMD-LNPs do not rely on successful genomic integration. Secondly, myo-AAVs need to successfully infect cells by evading immunological response and often result in production of neutralizing antibodies (Schulz et al., 2023). LNP immunogenicity is lower than that of AAVs and enables repeated dosing (Kenjo et al., 2021). Lastly, LNPs possess more tunable dynamics in terms of lipid composition and lipid molar ratios for targeted delivery to tissues (Vasileva et al., 2024).

Importantly, recent clinical results showed that LNPs were effective in restoring metabolic function in patients with propionic acidemia (Dolgin, 2024; Koeberl et al., 2024), demonstrating the utility of LNP platforms to restore cellular function in addition to use as a vaccine. Another LNP-based treatment, Onpattro®, is FDA approved and used clinically for treatment of hereditary ATTR amyloidosis (Adams et al., 2018). However, LNP’s are not without drawbacks entirely. One shortcoming of utilizing LNPs in the clinic would be the dosing regimen of EDMD1 patients. Dosing regimen will likely need to be first established in mouse studies to determine feasibility *in vivo* and allometric scaling for human doses has been accomplished for other LNP-based mRNA treatments (Attarwala et al., 2023). Clinical trials will then be necessary to determine LNP efficacy and toxicity in human treatment. Because emerin expression depleted over time, patients will likely require routine dosing, which will be patient-dependent or empirically determined. Despite this, improvement of patient outcomes and disease progression is paramount. Future work will also need to determine if treatment is most feasible with intramuscular injections where LNPs are to be targeted. LNPs can also be used for CRISPR-mediated restoration of emerin expression in muscle. The data presented within this study are a foundation for future studies utilizing EMD-LNPs for clinical applications with EDMD1 patients and potentially patients with other forms of EDMD.

