## Supplemental Figures for "Development of Emerin mRNA Lipid Nanoparticles to Rescue Myogenic Differentiation"

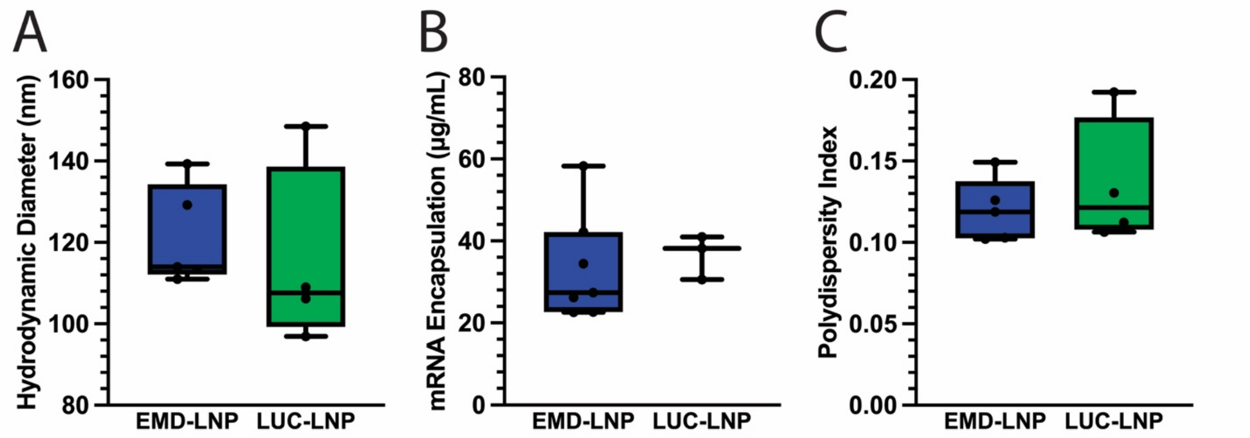


**Supplemental Figure 1. LNP formulation and characterization.** Characterization of emerin and luciferase mRNA LNPs including A, hydrodynamic diameter B, mRNA encapsulation amount, and C, polydispersity index.

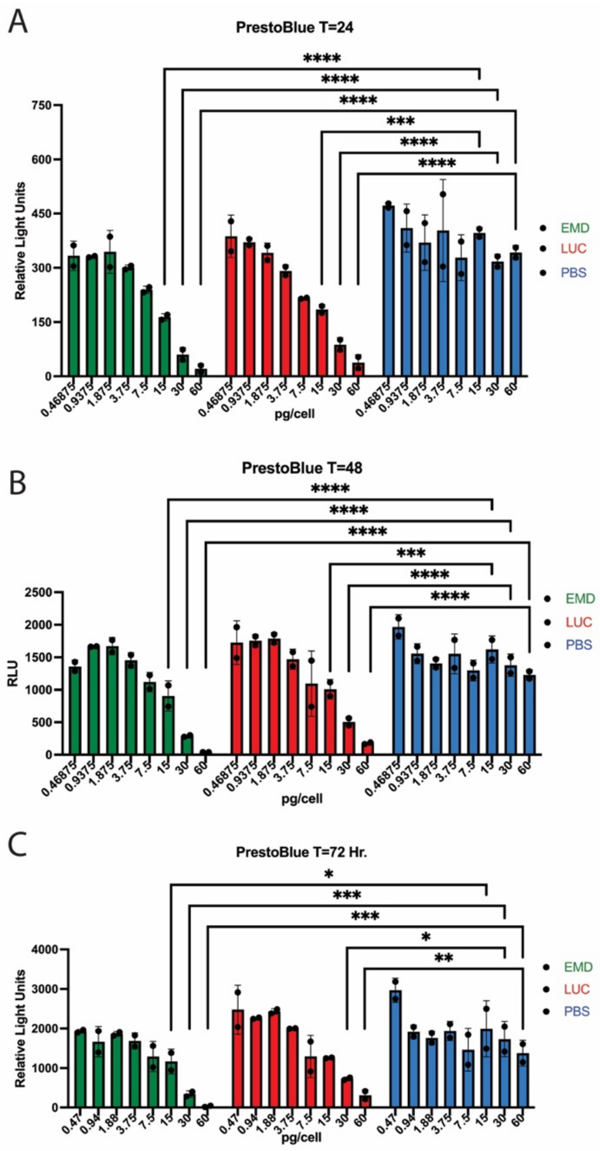


**Supplemental Figure 2. Full LNP dosing regimen used to monitor cell viability of emerin-null myogenic progenitors.** PrestoBlue cell viability assays were used to measure cellular metabolism in EMD$^{-/y}$ + PBS (PBS), EMD$^{-/y}$ + EMD-LNP (EMD LNP), or EMD$^{-/y}$ + luciferase mRNA LNP (LUC LNP) proliferating myogenic progenitors with increasing concentrations (0-60 pg/cell) of emerin or luciferase LNPs. Measurements were recorded every 24 hours for 72 hours. A, 24 hours after LNP incubation. B, 48 hours after LNP incubation. C, 72 hours after LNP incubation (N=2) (* p≤0.05; ** p≤0.01; *** p≤0.001)
